# RAPID: Real-time Animal Pattern re-Identification on edge Devices

**DOI:** 10.1101/2025.07.07.663143

**Authors:** András Zábó, Máté Nagy, Aamir Ahmad

## Abstract

Automatic re-identification of animals has significant potential for addressing pressing challenges of our times via population monitoring, individual health assessment or detailed behavior analysis. Although numerous computer vision based solutions have been proposed and many achieve high accuracy, those still remain unsuitable for real-time analysis and deployment on edge devices. Here, we introduce RAPID, an open-source^1^ algorithm for re-identifying patterned animal images at a rate exceeding 40-60 queries per second on standard PC or Laptop and over 10 queries per second on an inexpensive off-the-shelf edge device. RAPID operates efficiently in computationally restricted environments, relying solely on CPU, leaving GPU resources available for other tasks, all while maintaining state-ofthe-art accuracy. Our approach leverages SIFT descriptors, which continue to demonstrate competitive robustness and accuracy against recent traditional and deep-learning based methods. To counterpoise SIFT’s primary limitations its high-dimensional feature vectors and the associated computational costs - we incorporate recent advancements in vector similarity search and construct a database of feature vectors rather than database images, further accelerating the search process. We evaluate our method, RAPID, on public re-identification datasets; additionally propose a RAPID-based tool, FalseTagFinder, for cleaning benchmark dataset labels and as a demonstration, provide corrections for the StripeSpotter dataset.

## 1. Introduction

The ability to recognize individual animals among conspecifics was once exclusive to a handful of experts. However, with recent advancements in computer vision, this capability has begun to be shared with machines as well. Building upon the success of automated species recognition and diving one layer deeper, animal re-identification (reID) algorithms have been proposed to automate this challenging labor- and time-intensive task [1–5].

Image-based individual recognition algorithms primarily rely on image matching, wherein the most similar database image to the query is retrieved, and its corresponding identity is assigned to the processed image. Traditionally, the matching process consists of four main phases: (1) keypoint detection, where a detector identifies salient points in the query image; (2) keypoint description, which encodes the local environment of these points into descriptor vectors; (3) image similarity search within the database using these vectors; and (4) identity prediction based on the best-matching image. SIFT [6], SURF [7], and ORB [8] are the flagships of this classical approach, relying on handcrafted algorithms for detection and description. More recent handcrafted approaches [9, 10] have not gained wide popularity so far. In contrast, state-of-the-art methods leverage deep learning techniques, reaching beyond the classical approach. Convolutional neural network (CNN) based architectures now facilitate the extraction and description of local features, with some methods modifying the sequence of these two steps or even merging them into a single hybrid step [11–15]. Additionally, deep features higher-level, abstract image representations -have been introduced. These features, by their nature, cannot be localized at pixel level but capture image properties that are not accessible through traditional local features. Beyond the image feature extraction (phase 1 and 2), the similarity search part has also undergone significant advancements with the introduction of novel metric learning algorithms [16].

Both traditional and deep learning based methods, as well as hybrid approaches, have achieved high prediction accuracy on various reID datasets in recent years [17–21]. However limitations persist. Among traditional methods, SIFT remains one of the most robust algorithm, still competitive among today’s methods in case of patterned animals [22, 23], but its relatively large feature vector result in slower similarity search. Other algorithms may utilize lower dimensional vectors, which enable faster processing but sacrifice robustness as a trade-off. In deep learning based reID systems factors such as strong dependency on training data, abstract nature of deep features and high computational need are limiting in terms of both speed and edge deployment. It is worth to mention that the state-of-theart image matching algorithms [24–27] may seem to have a potential to make reID fast and robust, but matching sequential camera images at high frames per second (FPS) in robotic SLAM applications differs from matching images within large-scale databases. Beyond these limiting factors, the available datasets that contain individual identities are also still relatively scarce. Datasets available on LILA BC and Wildlife Datasets [22], besides often containing low resolution or blurred images, also lack viewpoint variability. Images are mainly taken from the ground, showing the animal only from the left or the right side. Datasets with a broader range of viewpoints could provide valuable information to support the development of novel approaches, leading to more robust and abstract individual representations.

Responding to these challenges, we present our work, which has multiple contributions to fostering research in animal reID. We chose SIFT features to ensure scale and rotational invariant descriptor vectors while preserving local keypoint information. Real-time processing, which we use as a synonym for 10 or higher FPS values, is achieved by shifting our focus to query feature vector matching, leveraging recent advancements in vector similarity search, rather than retrieving the most similar images from the database. Additionally, our traditional approach ensures RAPID remains lightweight compared to deep-learning based methods, also providing transparency throughout the pipeline without black-box components. The main contributions of this paper are as follows:

### 1. Real-time

RAPID runs real-time, it is able to process query images at a rate of 50 query images/second (averaged over 4 different datasets and two test hardware: Xeon E-2174G with 8 cores, 62 GB RAM and Ryzen 7 5700U with 16 cores, 16 GB RAM devices without GPU acceleration).

### 2. Scale and rotational invariant

Keypoints are extracted and described using the traditional SIFT algorithm, meaning that scale and rotational invariance is ensured. Furthermore keypoints can be localized on the images supporting further analysis in contrast to deep features.

### 3. Lightweight

RAPID can be deployed and efficiently run on edge devices. Leaving GPU resources free and relying only on CPU, it achieves 10 query image/second processing rate on a $100 off-the-shelf edge device (2× Cortex-A72) opening up the large variety of use cases and applications.

### 4. Benchmark corrections and FalseTagFinder

As part of our algorithm we provide a feature, FalseTagFinder, that identifies and highlights possible wrong labels within ground-truth datasets supporting researchers in database cleaning. As a demonstration, we also provide a cleaned version of the StripeSpotter dataset, where mislabels were detected with the help of FalseTagFinder.

## 2. Related work

Computer aided animal reID tools are being proposed for over a decade. First pioneering solutions were all built on local features, often requiring several minutes to process one single query image [30, 31]. These early methods relied on traditional feature extraction techniques, which are limited to detecting only local features. However, the past years’ deep learning revolution also left a mark in this field rapidly increasing the number of proposed methods [3, 17, 18, 33]. Unlike traditional approaches, deep learning based methods can extract not only local, but also deep features, offering potentially higher accuracy in challenging scenarios.

In this section, we provide a brief overview of previous studies utilizing local and deep features in the field of animal reID and present a high-level comparison of existing tools, focusing on local feature based solutions, as RAPID is a new candidate within this category.

### 2.1 Local features based solutions

Local feature based approaches represent images through precisely localizable keypoints and their described pixel environments. Traditional methods, which do not incorporate deep learning, are lightweight, transparent and have a plug-and-play nature, not needing training or exhaustive fine tuning.

Some of these algorithms such as I^3^S [30] and AmphIdent [31] are not suitable for real-time applications as they require minutes to process a single query image due to semi automated pipelines and need for human intervention. StripeSpotter [10] reports processing times of approximately 0.01 seconds per query, although this result may be misleading as it excludes feature extraction, and it is evaluated on small database sizes, less than 90 images, which in often unrealistic in real-life scenarios. In case of HotSpotter [32], an image is processed under 1-3 seconds. Its integration into multiple projects (IBEIS [34], Wild-Me [35], Conservation X Labs[36]) demonstrates its success but also led to maintenance being deprioritized, making installation and testing cumbersome. We note that these projects are not discussed as they primarily focus on database management and user-friendly GUIs rather than real-time query processing.

More recent algorithms utilize deep learning for local keypoint detection and description [33, 37]. As a comprehensive and gap-filler work, Wildlife-Tools [22], has been proposed recently. This modular Python package offers traditional SIFT, but also popular state-of-the-art deep learning based local feature extractors [38–40] and matchers [24, 26]. However, its primary goal is not to minimize query processing time but rather to serve as a general, userfriendly platform for animal reID. This flexibility and ease of use come at the expense of losing the ability to be highly optimized for speed.

### 2.2 Deep features based solutions

In case of deep feature based methods images are represented as high-dimensional embeddings rather than localized keypoints. These abstract vectors encode characteristics beyond pixel level information, capturing higher level properties of images. This seems to be particularly promising in case of open-set scenarios, where new individuals may appear in query that are not represented in the database.

Several species-specific models have been proposed, including those designed for gorillas [41], cetaceans [17], and cattle [42]. Additionally, the above mentioned WildlifeTools provides foundation models for general, multi-species re-identification. However, many of these methods do not report processing times [21], making it difficult to assess their real-time applicability. Despite their high accuracy in complex scenarios, deep feature based methods often require large-scale, diverse datasets for training and substantial computational resources.

### 2.3 Comparison of existing tools

In Tab. 1 we provide a high level overview of existing solutions. Two selection criteria were applied: utilizing local features and, preferably, being open-source, as our algorithm is a new candidate among such approaches. With the exception of WildlifeTools, all listed methods are traditional and do not incorporate deep learning based modules. However, WildlifeTools remains part of the comparison, as its framework allowed us to evaluate the state-of-theart deep learning based local feature extractors mentioned in Sec. 2.1. Additionally, we tested its MegaDescriptor-T-224 pretrained model to compare our algorithm with a deep feature based approach as well.

**Table 1.**
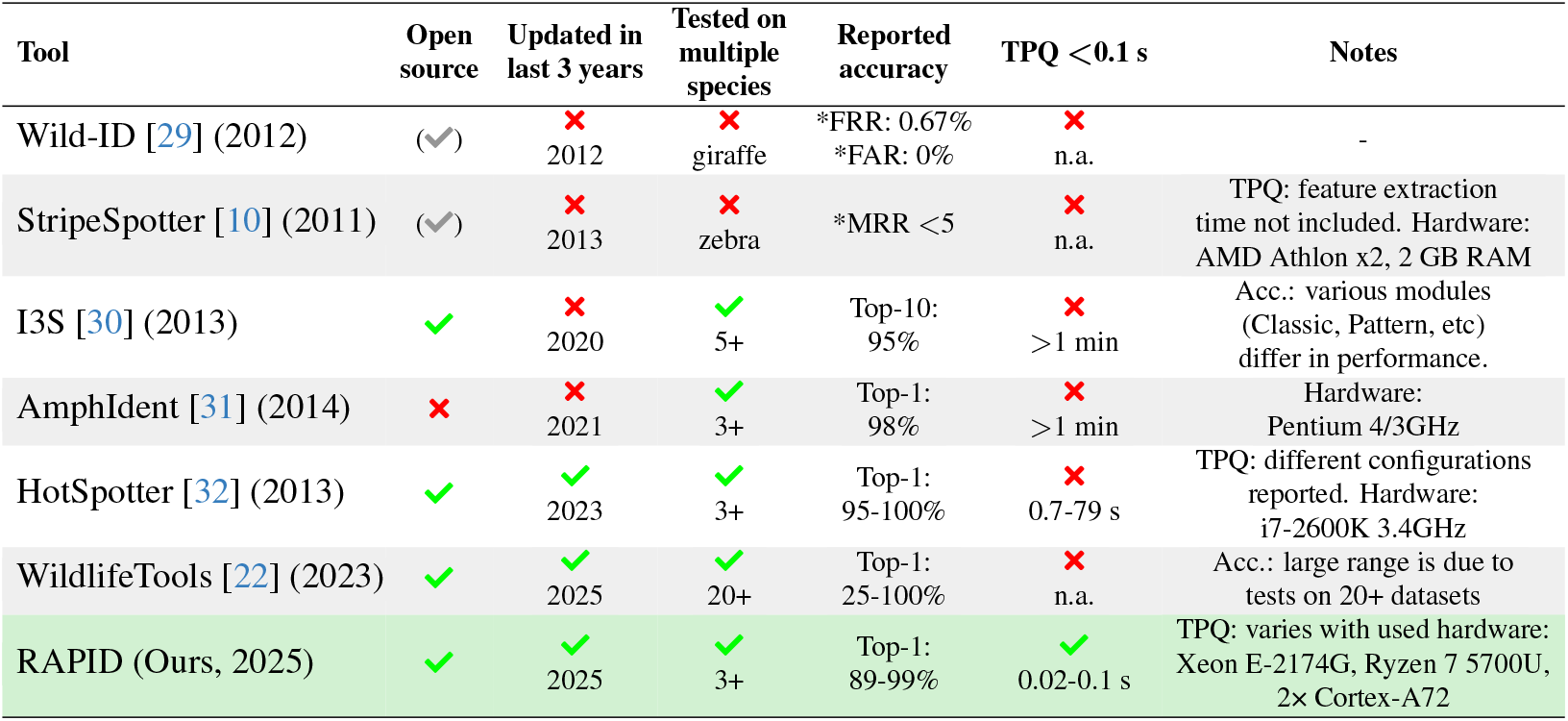
Existing local feature based reID tools. All values (except the last row) copied from related publications or GitHub pages. We use symbols 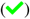 for fulfilled or not fulfilled 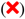 criteria. **(1) Tools:** Name/year of publication. **(2) Open source:** 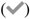 indicates open source tools, but not available any more on the provided links. **(3) Updated in last 3 years:** Provides information about last code maintenance/update. **(4) Tested on multiple species:** Species algorithm tested on. **(5) Reported accuracy:** Accuracy metrics provided by authors. Larger values corresponds to better performance, otherwise * indicates opposite behavior. FRR, FAR, MRR stand for False Rejection Rate, False Acceptance Rate and Mean Reciprocal Rank, respectively. Top-1 and Top-10 accuracy show the percentage of the retrieved correct ID among top 1 and top 10 predictions. **(6) TPQ *<*0.1 s:** Time per query (TPQ) values, n.a. if not reported. Cautious comparison is needed as publications refer to different processes executed under TPQ. **(7) Notes:** Further information for comparison.

| Tool | Open source | Updated in last 3 years | Tested on multiple species | Reported accuracy | TPQ < 0.1 s | Notes |
| --- | --- | --- | --- | --- | --- | --- |
| Wild-ID [29] (2012) | (✓) | ✗<br>2012 | ✗<br>giraffe | *FRR: 0.67%<br>*FAR: 0% | ✗<br>n.a. | - |
| StripeSpotter [10] (2011) | (✓) | ✗<br>2013 | ✗<br>zebra | *MRR < 5 | ✗<br>n.a. | TPQ: feature extraction time not included. Hardware: AMD Athlon x2, 2 GB RAM |
| I3S [30] (2013) | ✓ | ✗<br>2020 | ✓<br>5+ | Top-10: 95% | ✗<br>> 1 min | Acc.: various modules (Classic, Pattern, etc) differ in performance. |
| AmphIdent [31] (2014) | ✗ | ✗<br>2021 | ✓<br>3+ | Top-1: 98% | ✗<br>> 1 min | Hardware: Pentium 4/3GHz |
| HotSpotter [32] (2013) | ✓ | ✓<br>2023 | ✓<br>3+ | Top-1: 95-100% | ✗<br>0.7-79 s | TPQ: different configurations reported. Hardware: i7-2600K 3.4GHz |
| WildlifeTools [22] (2023) | ✓ | ✓<br>2025 | ✓<br>20+ | Top-1: 25-100% | ✗<br>n.a. | Acc.: large range is due to tests on 20+ datasets |
| RAPID (Ours, 2025) | ✓ | ✓<br>2025 | ✓<br>3+ | Top-1: 89-99% | ✓<br>0.02-0.1 s | TPQ: varies with used hardware: Xeon E-2174G, Ryzen 7 5700U, 2x Cortex-A72 |

## 3. Methodology

This section describes the structure of RAPID, which consists of two main components: offline and online. The offline component includes steps executed before query processing, while the online component means the real-time query processing itself. Additionally, several general modules are shared between both components. We begin with a high-level overview of RAPID’s pipeline, followed by the shared steps. We then detail the offline and online components. Finally, FalseTagFinder is elaborated.

### 3.1 High-level overview

RAPID takes a set of bounding box images (referred to simply as images from now on) of patterned animals with known IDs, forming the database image set. Each database image undergoes standardized preprocessing to ensure meaningful image comparisons. SIFT descriptors are then extracted and organized in an efficiently searchable tree-like structure. Additionally, database analysis is performed to support confidence score calculations. These last two steps constitute the offline component of RAPID.

In the online component, query images undergo the same preprocessing and features extraction as the database images. Once descriptor vectors are extracted from a query, nearest neighbor vectors and corresponding IDs are retrieved for each query vector. Retrieved IDs are weighted, and the ID with the highest weight is selected as RAPID’s prediction, accompanied by a confidence score.

### 3.2 General steps

#### 3.2.1 Image preprocessing

Both database and query images are processed the same way: they are resized to a width of 224 pixels while preserving the aspect ratio to prevent distortion of the animals’ patterns. The images are then converted into grayscale, and intensity values are normalized to ensures consistency in subsequent comparisons.

#### 3.2.2 Feature extraction

Feature extraction is also performed the same way for database and query images. The SIFT algorithm is used to detect and describe the top 150 most reliable keypoints in each image. SIFT was selected for four key reasons:

1. It produces more robust descriptor vectors compared to other traditional feature extractors such as SURF or ORB.
2. It is more lightweight than neural network based feature extractors, with faster and simpler setup and descriptor interpretation.
3. It still shows competitive accuracy among state-of-theart methods.
4. Our experiment showed that SIFT’s feature extraction is sufficiently fast, ensuring that it does not become a bottleneck for real-time performance.

### 3.3 Offline component

#### 3.3.1 Database building

Since SIFT features are extracted rapidly from the images, fast similarity search is the key to achieve real-time query processing. RAPID ensures this through three key design choices: (1) retrieving vectors associated with animal IDs during similarity search instead of database images, (2) decoupling database and query image processing, and (3) leveraging recent advancements in vector similarity search.

Unlike many reID tools that compute pairwise similarity between a query and all database image, requiring repeated processing of the database, RAPID extracts feature vectors from all database image once, and store them in an efficiently searchable tree-like structure, specifically in an ANNOY index. This eliminates the need for redundant database image processing, allowing feature extraction to be performed solely on the query image. ANNOY was chosen over other vector similarity search algorithms [43] due to its following properties:

1. Suitability for SIFT’s 128 dimensional descriptor vectors.
2. Efficiency for databases containing less than 1 million vectors (roughly 7,000 database images).
3. Low memory and CPU requirements, making it well-suited for edge deployment.

In addition to building the ANNOY index, a mapping is established between each database vector and its corresponding image and animal ID. This connection ensures that when a descriptor vector is retrieved, its source image and animal ID can be immediately identified.

We also note that previously built indexes can be reused and loaded into RAPID as well.

#### 3.3.2 Database analysis for confidence score calculation

To support query ID predictions with confidence scores, an analysis is performed on the database images. Since RAPID assigns weights to the predicted ID candidates, the ratio of the second-largest and the largest weight for a query can be calculated. First, these ratios are collected in cases where the predictions are correct (“True Match”), based on the ground truth labels. Next, to simulate incorrect predictions (“False Matches”), all descriptor vectors corresponding to the query animal are removed from the database index, and weight ratios are calculated again. The resulting distributions of True and False Matches, along with their corresponding Kernel Density Estimates (see Fig. 2), are used by the online component of RAPID to assign confidence scores to query weight ratios. The benefit of this approach is that the confidence scores will always depend on the database images, reflecting the variety of animals’ patterns, image qualities, etc.

**Figure 1.**
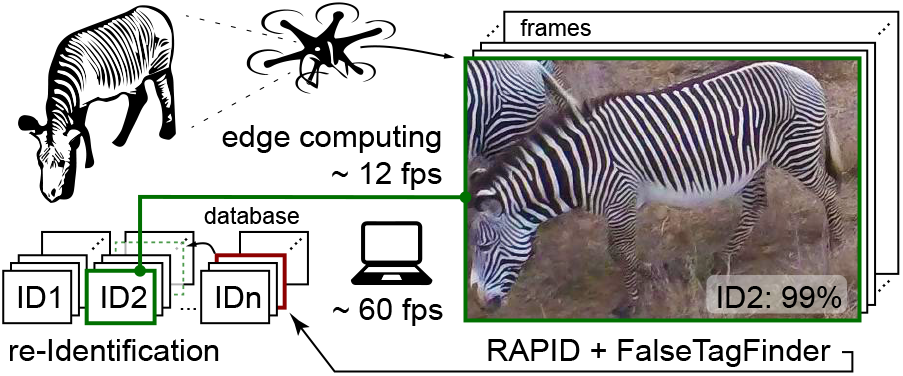
Overview of RAPID’s concept. RAPID predicts IDs, supported with confidence scores, to query animals at high FPS across diverse hardware and includes FalseTagFinder, a feature for database label cleaning.

**Figure 2.**
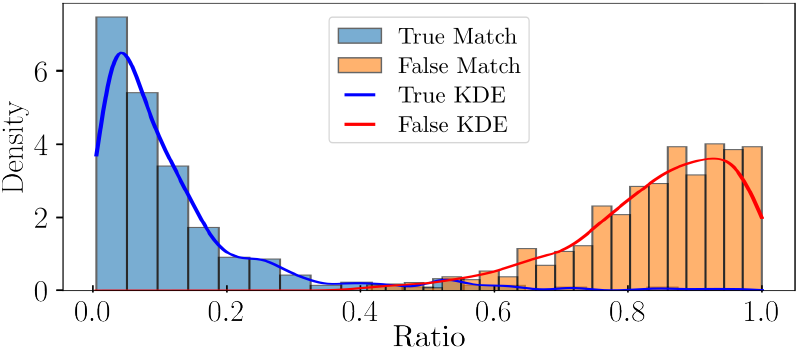
Weight ratio distributions of True and False Matches for AerialCattle2017. During query processing, retrieved IDs are ranked by their weights, and the ratio of the second-largest and the largest weights are calculated. Smaller ratio corresponds to more confident ID retrieval. Blue color shows the distribution of these ratios in case of correct (True Match), while orange in case of incorrect predictions (False Matches). Kernel Density Estimates are calculated to support later confidence score calculations.

### 3.4 Online component

#### 3.4.1 Similarity search

As queries went through the general image preprocessing and 150 most reliable keypoints were extracted and described, nearest neighbor database vectors are retrieved with corresponding Euclidean distances, animal and image IDs. Here we note that the popular Lowe’s ratio test [6] cannot be applied in our case as the same keypoint’s descriptor vector may appear multiple times in the database due to multiple images of the same animal.

#### 3.4.2 ID ranking

For all 150 matches of a query image, the retrieved IDs are weighted with the inverse of their corresponding nearest neighbor distance, resulting in larger weights for more similar vectors. If an ID appears multiple times among the matches, its weights are added up, further reinforcing the prominence of frequently occurring IDs. Finally, IDs are sorted by these final weights and the one in the first position serves as the prediction.

#### 3.4.3 Assigning confidence score

As an evaluation step, the ratio of the second-largest and the largest weights is computed, and the corresponding estimated density values are obtained from the Kernel Density Estimates. The confidence score is then assigned as the percentage of the True Match density relative to the sum of both density values. Overall, the smaller the weight ratio and the larger the confidence score, the more likely that the predicted ID is correct.

### 3.5 FalseTagFinder

Here, we summarize how RAPID can support researchers in cleaning database labels, a feature that we name FalseTagFinder. When using this feature of RAPID, the same steps are performed as elaborated above; however, in case of incorrect predictions—where the ground truth query ID does not match the algorithm’s prediction—an image is saved with relevant details. This assists researchers to investigate if the label is incorrect, however it is also a possibility that the algorithm has produced a false negative prediction. For an example of these outputs, see supplementary Fig. 6. The following information is provided in the saved images:

1. Query image and its image ID, animal ID and viewpoint information.
2. Six database images about the query animal, preferably with similar viewpoint. If less than six database images are available, some images will be shown multiple times.
3. Top three retrieved IDs are presented, each with two corresponding images. One image is the database image with the highest number of matching keypoints, while the other is the image with the single strongest matching keypoint.
4. Finally, a scatter plot is generated providing further information about the top 10 retrieved IDs, visualizing the number of retrieved source images and number of matching keypoints per IDs.

## 4. Experiments

In this section, we provide an exhaustive evaluation of RAPID. To gain a detailed picture of the capabilities and limitations of our algorithm we tested it with four public benchmark reID datasets accessed via LILA BC (Labeled Image Library of Alexandria: Biology and Conservation [28]) and Wildlife Datasets [22]. Furthermore, experiments were run on three different hardware, tracking both execution times and accuracy measures. Additionally, we demonstrate the use of FalseTagFinder by cleaning the StripeSpotter [10] dataset. First, the experimental setup is detailed, which is followed by a short summary about the used datasets and hardware platforms. Finally, evaluation metrics and results are provided.

### 4.1 Experimental setup

#### 4.1.1 Testing RAPID

For each dataset chosen for testing, we grouped the images by identities and randomly split these groups into five approximately equal-sized smaller subsets. Individuals with less than five images were excluded from the evaluation, as our focus was on the closed-set problem, where every query animal is also present within the database. Using these subsets, we performed a 5-fold cross-validation, ensuring that each animal image appeared in both the query and the database at least once during testing. In each fold, four sets (80% of the images) formed the database while the remaining one set (20% of the images) served as queries. To enhance variability and mitigate biases from a single random partitioning, this process was repeated three times per dataset yielding three independent 5-fold cross-validation tests. Results were averaged over these runs for each dataset and the whole process was conducted on all three hardware platforms.

#### 4.1.2 Testing various feature extractors

Within the framework of WildlifeTools we tested the query processing speed of various feature extractors: SIFT [6], SuperPoint [38], DISK [39], ALIKED [40] and also one deep feature based module, MegaDescriptor-T-224. In this case only PC and Laptop were used and only one set of query and database images were tested, since accuracy of these methods are known from [22], but no information is provided on algorithm speed.

#### 4.1.3 Testing FalseTagFinder

To demonstrate the use of FalseTagFinder, we cleaned the StripeSpotter dataset, which is known to contain some mislabels that have not yet been corrected. All images were included in the analysis and label adjustments were made iteratively based on the tool’s output until the results converged.

### 4.2 Datasets

For our experiments, we used four publicly available reID datasets: StripeSpotter [10], ATRW [44], AerialCattle2017[42] (referred to as AerialCattle) and GiraffeZebraID [45]. They contain images of Plains and Grevy’s zebras; Amur tigers; Holstein Friesian Cattle; and Plains zebras and Masai giraffes, respectively. All images were taken from the ground except in case of AerialCattle, where drone images introduced viewpoint variability to our tests. For all datasets, only individuals with at least five images were included in the tests. For StripeSpotter, only left side images, while for GiraffeZebraID, more challenging “frontleft” and “backleft” views were kept (more details in Supplementary Sec. 6). A general summary about test datasets are collected in Tab. 2.

**Table 2.**
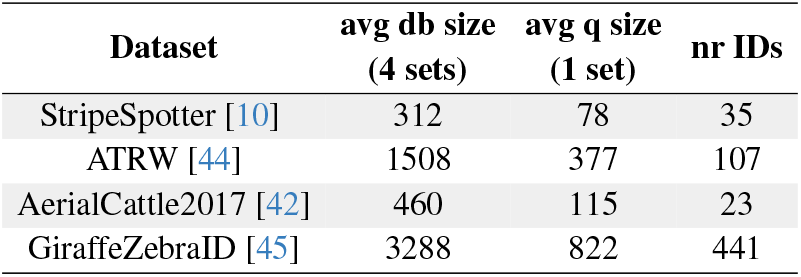
Datasets used for experiments. Individuals with at least five images were included in the tests. For StripeSpotter only left side, while for GiraffeZebraID more challenging “frontleft” and “backleft” images were kept. *Avg db size* and *avg q size* stand for number of images used in 5-fold cross-validation, *nr IDs* inform about the number of individuals.

| Dataset | avg db size<br>(4 sets) | avg q size<br>(1 set) | nr IDs |
| --- | --- | --- | --- |
| StripeSpotter [10] | 312 | 78 | 35 |
| ATRW [44] | 1508 | 377 | 107 |
| AerialCattle2017 [42] | 460 | 115 | 23 |
| GiraffeZebraID [45] | 3288 | 822 | 441 |

### 4.3 Hardware

Experiments were run on three different hardware: **(1) PC** - DELL Precision 3630 Tower with Intel(R)492 Xeon(R) E-2174G@ 3,80GHz CPU, 8 cores, 63 GB RAM; **(2) Laptop** - VivoBook-ASUSLaptop X513UA-M513UA with AMD Ryzen 7 5700U, Radeon Graphics, 16 cores, 16 GB RAM; and **(3) Edge** - Texas Instruments SK-TDA4VM with ARM Cortex-A72, 2 cores, 2.25 GB RAM. During experiments computers were not used for any other tasks. Furthermore, no CPU fine-tuning were applied, and GPU-s were not used.

### 4.4 Evaluation metrics

During execution we monitored the number of processed queries per seconds, referred to as FPS. Prediction performance was measured in form of Top-1 accuracy, which shows the percentage of correct predictions over queries. The common mean Average Precision (mAP) metric was not applicable in our case, as the algorithm returns identities not best matching image candidates.

### 4.5 Results

We present the results of RAPID with comparison to other tools obtained using various hardware while ensuring consistent test datasets and hardware configurations in all cases. First, we detail FPS performance of local feature extractors [6, 38–40] as implemented in WildlifeTools, along with a deep feature based approach using a pretrained model. Next, we show the speed and accuracy of our method RAPID. Finally, we demonstrate the reliability of RAPID’s confidence score, and database cleaning results achieved using RAPID’s FalseTagFinder.

Our evaluation of WildlifeTools demonstrated that realtime image processing is not possible within that framework (see Tab. 3). Query image processing using SIFT or recent state-of-the-art local feature descriptors (SuperPoint, DISK, ALIKED) all yielded FPS values well below 0.1. The highest performance, 6 FPS, was achieved on the PC using the AerialCattle dataset with the deep feature based MegaDescriptor-T-224 pretrained model. No significant variation was observed across the other three datasets or when tested on the Laptop (see Tab. 3 and Supplementary Tab. 5). Given the poor performance of WildlifeTools on the strongest hardware, we did not extended its testing to the edge device, as processing a single query image required between 0.5 to 5 minutes even on the PC.

**Table 3.** Query FPS on AerialCattle and StripeSpotter datasets with various feature extractors on two hardware. Evaluation on the same image set using WildlifeTools with its extractors, and RAPID with SIFT. MD-T-224 refers to deep feature based MegaDescriptor-T-224. RAPID, strongly outperforms compared methods.

| reID tool | feature extractor | AerialCattle |  | StripeSpotter |  |
| --- | --- | --- | --- | --- | --- |
|  |  | FPS (1/s) |  | FPS (1/s) |  |
|  |  | PC | Laptop | PC | Laptop |
| WildLifeT. | SIFT | <0.1 | <0.1 | <0.1 | <0.1 |
|  | SuperPoint | <0.1 | <0.1 | <0.1 | <0.1 |
|  | DISK | <0.1 | <0.1 | <0.1 | <0.1 |
|  | ALIKED | <0.1 | <0.1 | <0.1 | <0.1 |
|  | MD-T-224 | 6 | 5 | 5 | 4 |
| **RAPID (ours)** | **SIFT** | **59** | **54** | **38** | **37** |

To evaluate the performance of RAPID, FPS and Top-1 accuracy values were average over 15 runs for each benchmark datasets, based on three independent rounds of 5-fold cross-validations (Tab. 4, SIFT column). The results demonstrate the superior speed of RAPID, achieving 40-60 FPS across various species on both PC and Laptop. On the edge device, RAPID maintained 10 FPS — approximately twice the speed of WildlifeTools’ best performance on the most powerful hardware. Regarding accuracy, RAPID achieved 89%, 95%, 99% and 99% Top-1 accuracy on the tested bechmark datasets.

**Table 4.**
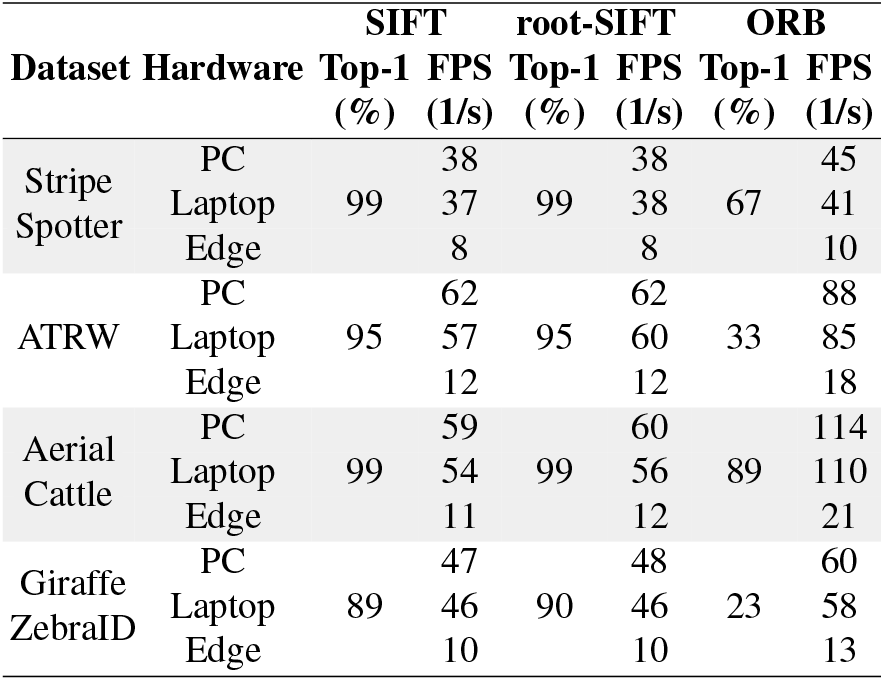
Accuracy and speed of RAPID for multiple datasets, hardware and descriptors.

In addition we conducted the same experiments with RAPID, but using ORB instead of SIFT and tested root-SIFT descriptor vectors as well (Tab. 4, root-SIFT, ORB columns). As expected, RAPID with ORB provided higher FPS values but at a cost of reduced accuracy. Consistent with [32] but in contrast to [46] we found no significant change in accuracy when using root-SIFT vectors instead of the classical ones (only in case of GiraffeZebraID the accuracy increased from 89% to 90%, otherwise stayed the same).

We also tested how the confidence score changes with varying query animal scenes. Here, we provide an example from a video where an animal passes behind another (Fig. 3). We tracked the confidence score and previously discussed weight ratio during sequential frames. Our results showed that prediction confidence drops down when 2 animals are present in the bounding box at the same time, while it increases as one occupies most of the image.

**Figure 3.**
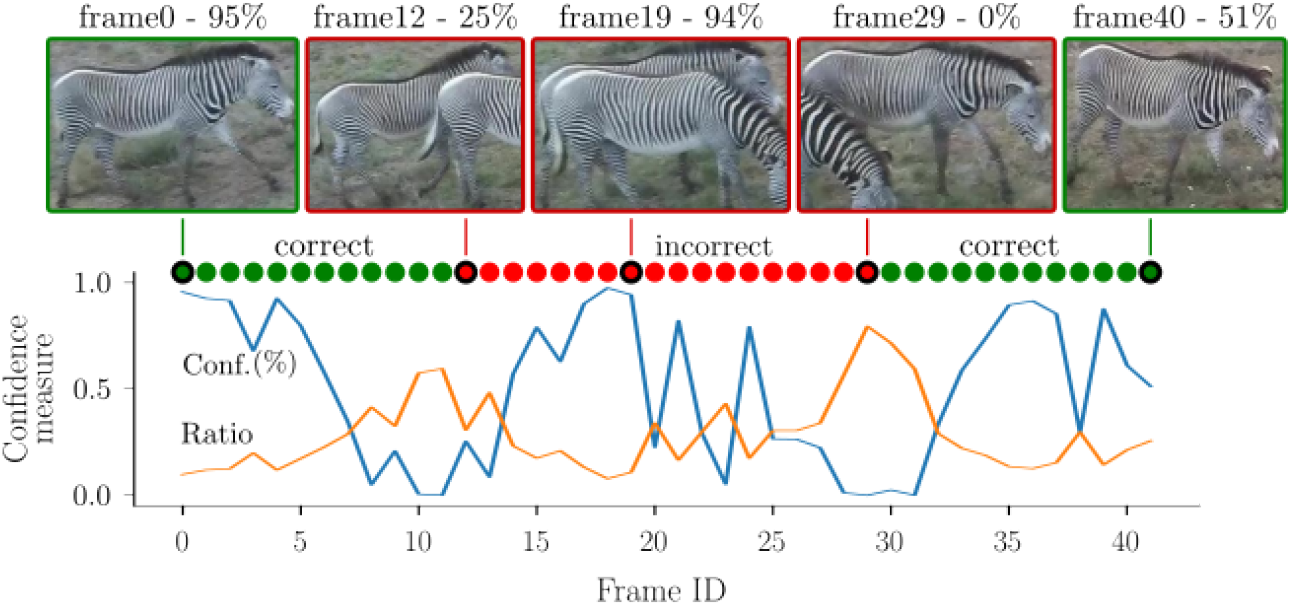
Performance of the confidence score. Tracking an individual over 40 frames and plotting confidence values for each frame. The blue line shows how the confidence score drops down as two animals are within the bounding box, and it also increases as it correctly detects the animal in front. However, predictions are incorrect in this case (red circles at the top) as the bounding box has the ID of the animal passing behind. The yellow line shows how the ID weight ratio changes in time, indicating an inverse behavior to the confidence score. The smaller the ratio, the more confident the prediction is.

**Figure 4.**
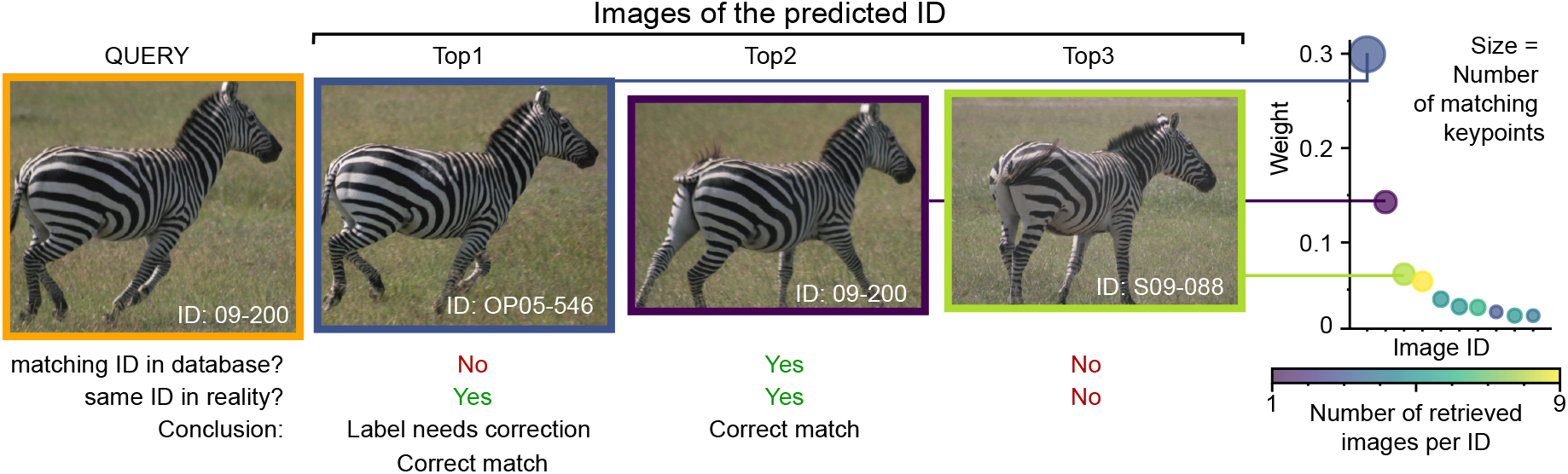
Label correction supported by FalseTagFinder. Output shows the query image and the three images with most keypoint matches corresponding to the three most confident ID predictions. Based on current labels, the correct ID appears in the second position. Beyond RAPID’s suggestion, careful pattern analysis and viewpoint similarities further reinforce the probably incorrect label in the original dataset for Top1 prediction, and this shall be corrected. Scatter plot (on the right side) shows additional information for top10 predicted IDs, to provide further properties of the prediction process.

Finally, we present how our tool’s FalseTagFinder feature can be used to clean datasets by demonstrating on a case study, the StripeSpotter database. Based on the suggestions provide by the tool, we identified and corrected various errors: side information was mislabeled in five instances; IDs from the existing list of individuals were assigned to previously unlabeled images in three cases; individuals were missing from the list of IDs (previously those images were labeled incorrectly to a different animal’s ID,and we solved this by introducing new IDs) in four cases; and images were mislabeled to different individuals in 20 instances. We provide the list of corrections in the Supplementary materials.

## 5. Discussion

Our results showed that RAPID has superior speed over existing methods while relying solely on CPU. Despite using SIFT, which is commonly considered not suitable for real-time applications, we achieved 40-60 FPS during query processing on a Laptop and PC. Furthermore 10 FPS was achieved on an off-the-self edge-device, finetuning CPU performance could result in even higher FPS. To the best of our knowledge this is the first work providing real-time query processing and optimal deployment on edge devices. We also showed that accuracy was not sacrificed for speed, achieving similar Top-1 accuracy results as other state-ofthe art methods. However, RAPID with ORB features (instead of SIFT) achieved 100+ FPS by trading off accuracy significantly on average. Interestingly, in case of AerialCattle, when FPS exceeded 100, accuracy of RAPID with ORB was suprisingly high, 89%. This suggests that in some cases, if the pattern type and images allow, it may be worth using ORB instead of SIFT. We note that Wildlife-Tools achieved low FPS values, since for local features only pairwise matching is currently available. In the case of MegaDescriptor-T-224, the matching happens based on another strategy, thus resulting as expected in the best performance among all tested methods within the framework. However, even as compared to this, RAPID showed 10x better FPS values. We also note that using only CPU (instead of GPU) may also contribute to the inferior query processing times of WildlifeTools. In contrast to several existing methods, RAPID’s further benefit is that it takes raw bounding boxes, and no further image manipulation is 8 required. This freedom and its CPU-based approach result in large potential for RAPID to be used as a modular tool that can be incorporated into other pipelines. RAPID’s feature to provide confidence scores could also be crucial for real-life applications that use real-time identification. As animals partially occlude each other, or in other cases when multiple individuals are present simultaneously in the same bounding box, the confidence of prediction drops. This may serve as a main characteristic to build upon for further reID related applications. Finally, we demonstrated that FalseTagFinder feature of RAPID is suitable for supporting and speeding up database cleaning.

Despite the promising results, RAPID and related features have several limitations. First of all, RAPID is only suitable for patterned animals and works in a “closed-set” case where the task is to find the ID of an individual from an existing database. This may be suitable for many, but not all, real-life applications. Furthermore, if an individual is represented by very few images within the database, or no similar viewpoint is available there as in the query, the prediction is likely to fail. This was our experience when cleaning StripeSpotter database. FalseTagFinder is not fully automated (with its pros and cons) and requires human intervention and decision-making to change the flagged labels. For certain cases, where the database does not contain a large enough number or variation in images per ID, FalseTagFinder may provide “False Negative” flags (for images that were correct but resulted in a poor match typically due to different view points), and thus this may slow down the cleaning process. However, the whole procedure is still faster overall than without the support of RAPID.

## 6. Conclusion

In this paper, we presented RAPID, a real-time, lightweight SIFT-based reID algorithm designed for patterned animals.Additionally, we introduced its FalseTagFinder feature for cleaning ground truth labels, along with proposed corrections for the publicly available StripeSpotter dataset. To the best of our knowledge, this is the first work achieving realtime query processing and edge deployment in this context. RAPID’s efficiency, low computational cost, and ability to provide confidence scores make it well-suited for a range of applications, from research to population monitoring in animal husbandry and wildlife.

Future work should address open-set re-identification challenges and enhancing FalseTagFinder to classify labeling error types and provide more interpretable outputs.

We believe that RAPID could significantly improve automated data collection on the individual level. At a broader scale, population-level analysis enabled by these methods could support wildlife conservation and nature protection.

