## Supplementary information for "RAPID: Real-time Animal Pattern re-Identification on edge Devices"

### Supplementary Material

### Supplementary Methods

#### Preparation step for using RAPID

Once RAPID is successfully installed, only two sets of images are required to execute the algorithm. One set, consisting of images with known IDs serves as the database, and the remaining images with unknown labels form the query set. Database images must be named following the format *animalID_side_imgID.imgtype* (*e.g. tiger3_front-left_23.jpg* or *zebra8_nosidesinfo_214.png*). These images should be already cropped bounding boxes.

#### Datasets

For algorithm development and early-stage testing we selected the StripeSpotter dataset [10], which, despite containing some unreliable labels, offers favorable properties. This dataset contains 824 images of 46 individuals, Plains and Grevy's zebras. Images were taken from the ground, showing animals either from the left or right side. Additionally, only a small number of bounding boxes contain multiple individuals, making this dataset suitable for isolating algorithmic challenges from those arising due to complex input images. We ran tests separately for the images showing the right and the left sides. We performed the deeper analysis only on one of these datasets, showing the left side, as that contains more individuals after filtering (a total of 35). We used whole bounding boxes, as opposed to the cropped bounding boxes (which are also part of the dataset) showing only the flanks of the animals without any background or other distracting features. After filtering for minimum number of images and left side photos, 35 individuals remained for testing (in contrast to 30 for right side). The average set size was 78 images, resulting in queries with this size and database sizes 4 times as large.

To evaluate how the method generalizes, species other than zebras were also tested. We run experiments on the ATRW dataset [44]. It consists of 3393 images of 274 Amur tigers, extracted from video footage taken from the ground. However, labels are provided only for 1886 images with 107 animal IDs. Additional challenge beyond moving to another species was that in contrast to StripeSpotter dataset, here the annotations do not provide information about viewpoints resulting in mixed side views. Filtering on minimum number of images per individuals resulted in 107 remaining IDs with an average set size of 377 images.

To further examine how accuracy changes with different viewpoints, we conducted tests on the AerialCattle2017 dataset [42]. These images were taken from drones, capturing various tracks of 23 Holstein Friesian Cattle. Large number, 46,340 images form this dataset as the continuous frames of several video footages. In order to avoid highly similar images, we randomly picked 575 images, 25 per each 23 individuals for testing, resulting in 115 images query size and four times larger databases.

As a final test, we ran experiments on the GiraffeZebraID dataset [45] which contain ground images from various viewpoint about 2 species, Plains zebras and Masai giraffes. By this, we gained further insight into how well the algorithm separates different pattern types. Since the rich annotation information provides viewpoint labels as well, we filtered for "left" side, but kept more challenging images with "frontleft" or "backleft" annotations, too. After these steps 441 individuals remained for testing with average set size of 822 images.

| reID tool | feature extractor | ATRW FPS (1/s) | | GiraffeZebraID FPS (1/s) | |
|---|---|---|---|---|---|
| | | PC | Laptop | PC | Laptop |
| WildLifeT. | SIFT | <0.1 | <0.1 | <0.1 | <0.1 |
| | SuperPoint | <0.1 | <0.1 | <0.1 | <0.1 |
| | DISK | <0.1 | <0.1 | <0.1 | <0.1 |
| | ALIKED | <0.1 | <0.1 | <0.1 | <0.1 |
| | MD-T-224 | 6 | 4 | 5 | 4 |
| **RAPID (ours)** | **SIFT** | **62** | **57** | **47** | **46** |

Table 5. **Query FPS on ATRW and GiraffeZebraID datasets with various feature extractors on two hardware.** Evaluation on the same image set using WildlifeTools with its extractors, and RAPID with SIFT. MD-T-224 refers to deep feature based MegaDescriptor-T-224. RAPID, strongly outperforms compared methods.

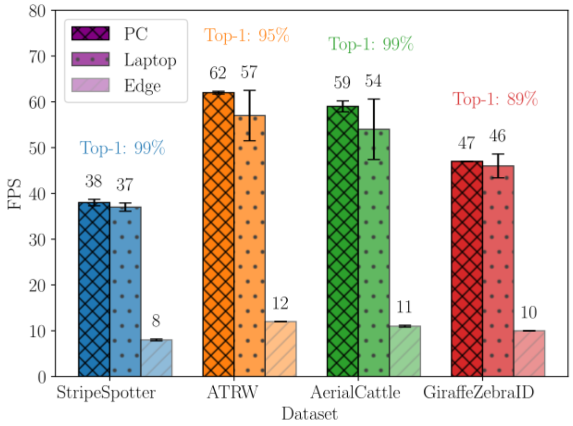


Figure 5. **Performance of RAPID with SIFT**.

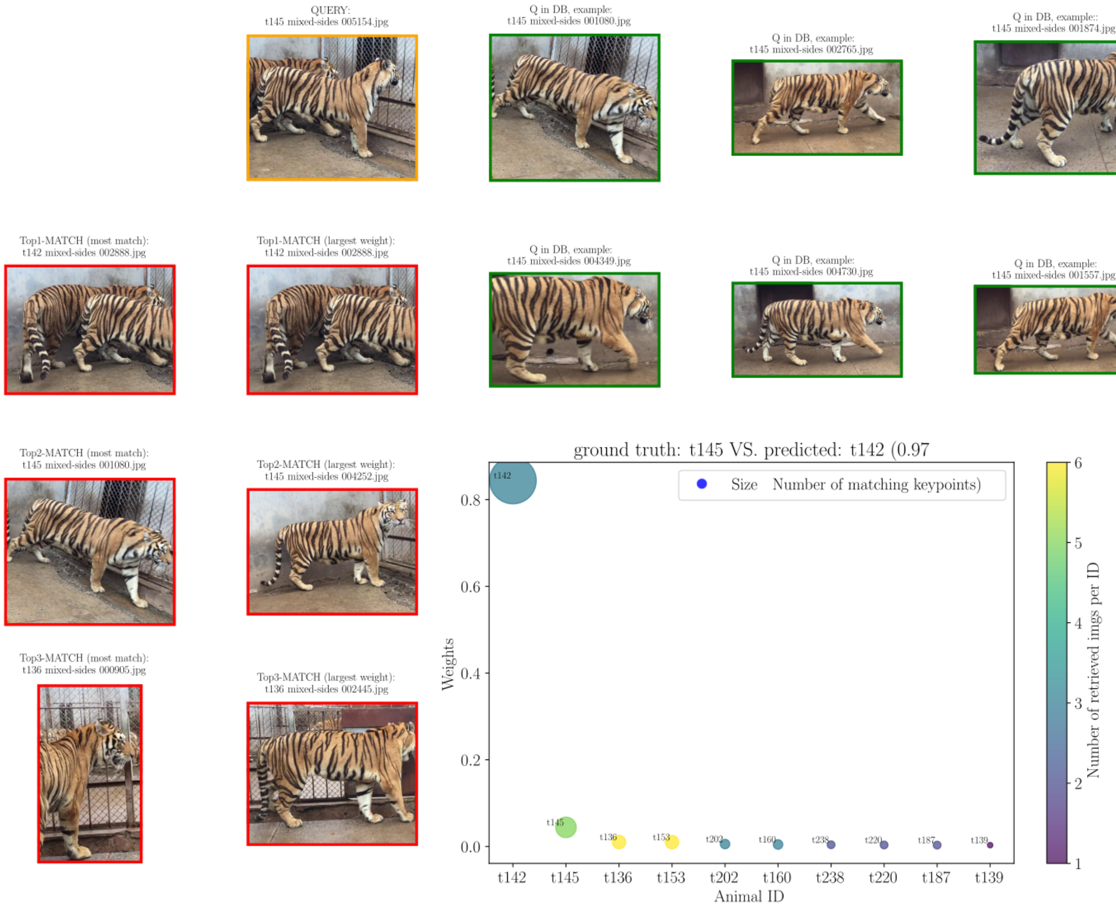


Figure 6. **Raw output of FalseTagFinder**. Query animal is shown in orange, corresponding six database images in green frame, possibly with matching side annotation (*e.g*. left or right). If the query animal is represented with less than six images, some of them will appear multiple times. Three most confident retrieved IDs and corresponding two images for each are shown with red image border. Scatter plot shows further information about the top 10 retrieved IDs.

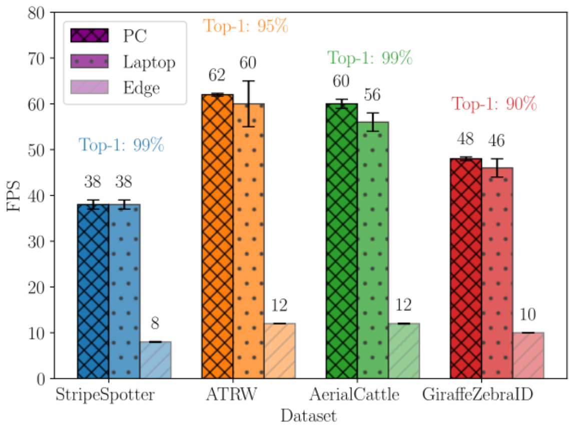


Figure 7. **Performance of RAPID with rootSIFT**.

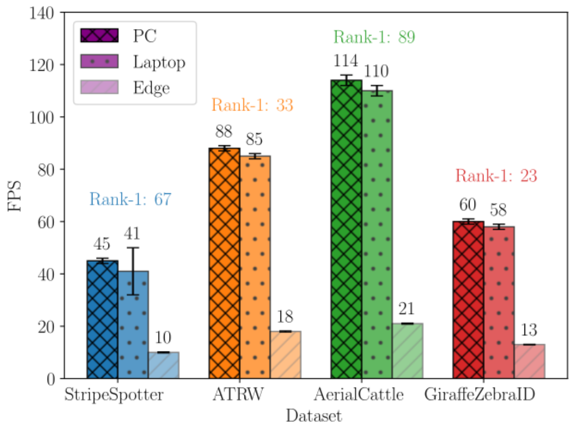


Figure 8. **Performance of RAPID with ORB**.
